# Updating the 97% identity threshold for 16S ribosomal RNA OTUs

**DOI:** 10.1101/192211

**Authors:** Robert C. Edgar

## Abstract

The 16S ribosomal RNA (rRNA) gene is widely used to survey microbial communities. Sequences are often clustered into Operational Taxonomic Units (OTUs) as proxies for species. The canonical clustering threshold is 97% identity, which was proposed in 1994 when few 16S rRNA sequences were available, motivating a reassessment on current data. Using a large set of high-quality 16S rRNA sequences from finished genomes, I assessed the correspondence of OTUs to species for five representative clustering algorithms using four accuracy metrics. All algorithms had comparable accuracy when tuned to a given metric. Optimal identity thresholds that best approximated species were ∼99% for full-length sequences and ∼100% for the V4 hypervariable region.

## Background

Next-generation sequencing of the 16S ribosomal RNA (rRNA) gene has revolutionized the study of microbial communities in environments ranging from the human body (Cho and Blaser, 2012; Pflughoeft and Versalovic, 2012) to oceans (Moran, 2015) and soils (Hartmann *et al.*, 2014). Data analysis in such studies typically assigns 16S rRNA sequences to Operational Taxonomic Units (OTUs). Many OTU clustering methods have been proposed (for example (Seguritan and Rohwer, 2001; Schloss and Handelsman, 2005; Schloss *et al.*, 2009; Ye, 2011; Edgar, 2013; Rideout *et al.*, 2014)), most of which use a threshold of 97% sequence identity. Typically, this threshold is considered given rather than as a tunable parameter, following the conventional wisdom that 97% corresponds approximately to species (Seguritan and Rohwer, 2001; Schloss and Handelsman, 2005; Westcott and Schloss, 2017). The 97% threshold was proposed in 1994 (Stackebrandt and Goebel, 1994) when few 16S rRNA sequences were available, raising the question of whether this value is supported by the much larger datasets currently available. In this work, I used a high-quality set of 16S rRNA sequences from known species to investigate whether the 97% threshold is a good approximation to species, whether a better threshold can be identified, and whether clustering algorithms can be ranked by quality.

OTU clustering is most commonly used in analysis of next-generation amplicon reads of the 16S rRNA gene. These reads have errors due to PCR and sequencing which can cause large numbers of spurious OTUs (e.g., (Huse *et al.*, 2010; Edgar and Flyvbjerg, 2014)). Thus, in practice, low OTU quality may be due to inadequate error filtering rather than the clustering algorithm. Here, I focus on OTUs of correct sequences to investigate whether algorithms differ in their ability to reproduce species classifications by taxonomists. While it could be of interest to investigate the tolerance of clustering algorithms to errors, this is a complex issue beyond the scope of the work reported here. Also, state-of-the-art denoisers have been shown to accurately recover biological sequences from 454 and Illumina amplicon reads (Quince *et al.*, 2009; Callahan *et al.*, 2016; Edgar, 2016) suggesting that the best strategy for amplicon reads is to cluster denoised sequences, in which case the clustering problem is well-modeled by error-free sequences from known species. Several OTU quality metrics have been proposed, including richness (e.g., (Sun *et al.*, 2009)), normalized mutual information (Cai and Sun, 2011; Zheng *et al.*, 2012) and Matthews’ Correlation Coefficient (Schloss and Westcott, 2011). I investigated whether different quality metrics give consistent algorithm rankings, which would support published claims that some algorithms generate objectively superior OTUs (e.g., (Cai and Sun, 2011; Schloss, 2008; Schloss and Westcott, 2011; Westcott and Schloss, 2017)).

## Methods

### HiQFL and HiQV4 databases

I constructed a database (HiQFL) of high-quality, full-length 16S rRNA sequences from known species as follows. I downloaded all prokaryotic genome assemblies from GenBank (Benson *et al.*, 2013) that were annotated as “Complete” in the in the *assembly_summary_ genbank.txt* file on Feb 15th, 2017. 16S rRNA gene sequences were identified using SEARCH_16S (Edgar, 2017). If any wildcard letters or ambiguity codes were found in a 16S rRNA sequence, all sequences from its assembly were discarded to avoid ambiguous sequence identities and ensure that intra-genome variation between 16S rRNA paralogs was accurately represented. One copy of each identical sequence from each assembly was retained. HiQFL contains 16 741 sequences from 6 240 assemblies of 2 512 species. Some species have many assemblies, with most for *Escherichia coli* (1 115 assemblies) and *Salmonella enterica* (1 035), while 1 106 species have exactly one assembly. To create a dataset with less taxonomic bias, I created the HiQFL_1 database by selecting one assembly at random for each species. The V4 hyper-variable region is currently a popular target for next-generation sequencing. To test clustering on high-quality V4 data, I constructed the HiQV4 and HiQV4_1 databases by extracting the segment between the primers V4F=GTGCCAGCMGCCGCGGTAA and V4R=GGACTACHVGGGTWTCTAAT (Kozich *et al.*, 2013)from HiQFL and HiQFL_1 respectively.

### OTU quality metrics

I used four quality metrics *RR*, *NMI*, *MCC*_sp_ and *Bij*. Richness ratio (*RR*) is min(*S*, *N*)/max(*S*, *N*) where *S* is the number of species and *N* is the number of OTUs.

Normalized mutual information (*NMI*) (Cover and Thomas, 1991) is an information theory measure of the mutual dependence between two frequency distributions. Matthews’ Correlation Coefficient (*MCC*) (Matthews, 1975; Baldi *et al.*, 2000) measures the accuracy of a binary classifier as a correlation between predicted and known values. I defined the correlation between OTUs and species (*MCC*_sp_), by considering a pair of sequences to be correctly classified if they belong to the same species and are in the same OTU. This differs from the metric (*MCC*_SW_) of (Schloss and Westcott, 2011) where a pair is considered to be correctly classified if they have ≥97% identity and are in the same OTU (see Discussion). I defined bijection (*Bij*) as the fraction of species that have 1:1 correspondence with an OTU. All four metrics have a maximum value of one indicating the best possible quality. For further details and discussion, see the Supplementary Material.

### Clustering algorithms

I tested the following clustering algorithms: nearest-neighbor (*NN*, also called single-linkage), average-neighbor (*AN*, also known as UPGMA or average-linkage), furthest-neighbor (*FN*, also called complete-linkage), OptiClust (*OC*) (Westcott and Schloss, 2017), and abundance-sorted greedy clustering (*AGC*) (Ye, 2011). For NN, AN, FN and OC I used mothur v1.39.5 (Schloss *et al.*, 2009) (commands given in Supplementary Files). I implemented AGC in a Python script that accepts a mothur distance matrix as input to ensure that that the same identities were used by all algorithms.

### Optimal thresholds

I ran each clustering algorithm on the four HiQ databases with thresholds ranging from 96% to 100% in steps of 0.1%. For each database, clustering algorithm and quality metric, I identified the optimal threshold as the tested threshold which gave the largest value of the metric.

### Conspecific probability

It is well-known that some pairs of species have 16S rRNA sequences with >97% identity and using a fixed threshold cannot reliably identify species (e.g., (Schloss, 2010)). This correspondence between identity and species can be investigated independently of clustering by measuring the probability that two sequences are conspecific (i.e., belong to the same species) as a function of identity. Let the *conspecific probability P_cs_*(*D* | *d*(*X*, *Y*)) be the probability that two sequences *X*, *Y* selected at random from a distribution *D* belong to the same species given a measure *d* of pair-wise distance between *X* and *Y*. I calculated *P*_*cs*_ for each HiQ database by assuming that sequences are drawn at random from the database with equal probabilities. Pair-wise identities calculated by mothur were binned into intervals of 0.5%. For each bin (e.g. 97.0%≤*d*<97.5%), let *M*_*d*_ be the total number of pairs and *m*_*d*_ be the number of pairs which belong to the same species, then *P*_*cs*_(*d*) = *m*_*d*_ /*M*_*d*_.

### Assessment of pair-wise distances

Mothur distances are calculated from a multiple alignment constructed using an algorithm based on the NAST strategy (DeSantis *et al.*, 2006) which introduces misalignments to preserve a fixed number of columns. I compared alignments by mothur and CLUSTALW v2.1 (Thompson *et al.*, 2002) on 16S rRNA sequences from (Kozich *et al.*, 2013). I took the 100 most abundant unique sequences in the reads assigned to soil samples (*soil100*) and constructed alignments for all pairs. For each pair-wise alignment, I calculated identity as the number of columns containing identical letters divided by the number of columns containing at least one letter.

### Adverse triplets

It has been proposed (Schloss and Westcott, 2011; Westcott and Schloss, 2017) that OTUs should be constructed such that all pairs of sequences with identity ≥97% are assigned to the same OTU and all pairs <97% are assigned to different OTUs. This constraint cannot be satisfied if there is an *adverse triplet* {A, B, C} with pair-wise distances A-B ≥97%, B-C ≥97% and A-C <97% because A-B and B-C imply that A, B and C should be assigned to the same OTU while A-C implies that A and C should be assigned to different OTUs. If a solution exists, an adverse triplet cannot be present because all pair-wise constraints are satisfied. Therefore, a solution exists if, and only if, there are no adverse triplets in the data. To investigate whether this is an issue in practice, I identified adverse triplets of species in the HiQ16_1, HiQV4_1 and soil100 datasets using the mothur distance matrixes.

## Results

### Optimal thresholds

Optimal thresholds are given in Table 1; metric values for all thresholds are given in the Supplementary Files. All algorithms achieve comparable maximum scores with all metrics. No algorithm is consistently better than any other, showing that algorithms cannot be meaningfully ranked by OTU quality. Optimal thresholds are all higher than 97%, especially on V4 where the optimal threshold is 100% for 9/20 of algorithm-metric combinations on HiQV4 and 17/20 on HiQV4_1.

**Table 1.** Optimal thresholds and metric values. Metric values for all thresholds are provided in the supplementary data.

| Data | Metric | Maximum metric |  |  |  |  | Optimal identity threshold |  |  |  |  |
| --- | --- | --- | --- | --- | --- | --- | --- | --- | --- | --- | --- |
|  |  | NN | AN | OC | FN | AGC | NN | AN | OC | FN | AGC |
| HiQFL | RR | 0.99 | 0.99 | 0.99 | 1.00 | 0.98 | 99.3 | 99.1 | 99.0 | 98.8 | 99.1 |
|  | MCC<sub>sp</sub> | 0.86 | 0.86 | 0.85 | 0.85 | 0.85 | 99.3 | 98.5 | 98.4 | 97.8 | 98.0 |
|  | Bij | 0.62 | 0.63 | 0.62 | 0.62 | 0.63 | 99.4 | 99.4 | 99.4 | 99.3 | 99.4 |
|  | NMI | 0.95 | 0.95 | 0.95 | 0.95 | 0.94 | 99.3 | 98.8 | 98.9 | 97.8 | 98.0 |
| HiQFL_1 | RR | 1.00 | 0.98 | 0.98 | 0.99 | 0.99 | 99.5 | 99.4 | 99.4 | 99.2 | 99.3 |
|  | MCC<sub>sp</sub> | 0.55 | 0.61 | 0.61 | 0.64 | 0.59 | 99.6 | 99.4 | 99.4 | 99.0 | 99.3 |
|  | Bij | 0.68 | 0.68 | 0.67 | 0.67 | 0.67 | 99.5 | 99.4 | 99.4 | 99.4 | 99.4 |
|  | NMI | 0.96 | 0.97 | 0.97 | 0.97 | 0.97 | 99.5 | 99.2 | 99.4 | 99.1 | 99.3 |
| HiQV4 | RR | 0.85 | 0.97 | 0.92 | 0.98 | 0.94 | 100.0 | 99.6 | 99.6 | 99.6 | 99.6 |
|  | MCC<sub>sp</sub> | 0.78 | 0.79 | 0.79 | 0.79 | 0.78 | 100.0 | 99.6 | 99.6 | 98.8 | 99.3 |
|  | Bij | 0.51 | 0.51 | 0.51 | 0.51 | 0.52 | 100.0 | 100.0 | 100.0 | 100.0 | 100.0 |
|  | NMI | 0.93 | 0.93 | 0.93 | 0.93 | 0.93 | 100.0 | 99.6 | 99.6 | 99.2 | 100.0 |
| HiQV4_1 | RR | 0.93 | 0.93 | 0.93 | 0.93 | 0.93 | 100.0 | 100.0 | 100.0 | 100.0 | 100.0 |
|  | MCC<sub>sp</sub> | 0.48 | 0.50 | 0.50 | 0.49 | 0.48 | 100.0 | 99.6 | 99.3 | 99.6 | 100.0 |
|  | Bij | 0.56 | 0.56 | 0.56 | 0.56 | 0.56 | 100.0 | 100.0 | 100.0 | 100.0 | 100.0 |
|  | NMI | 0.95 | 0.95 | 0.95 | 0.95 | 0.95 | 100.0 | 100.0 | 100.0 | 100.0 | 100.0 |

### Conspecific probabilities

See Fig. 1; numerical values are given in Supplementary Table S1. Conspecific probabilities for the four databases are quite different, illustrating that the probability depends on the gene segment (full-length or V4) and on the distribution from which sequences are sampled; i.e., on the composition and abundance distribution of species in the data.

**Fig. 1.**
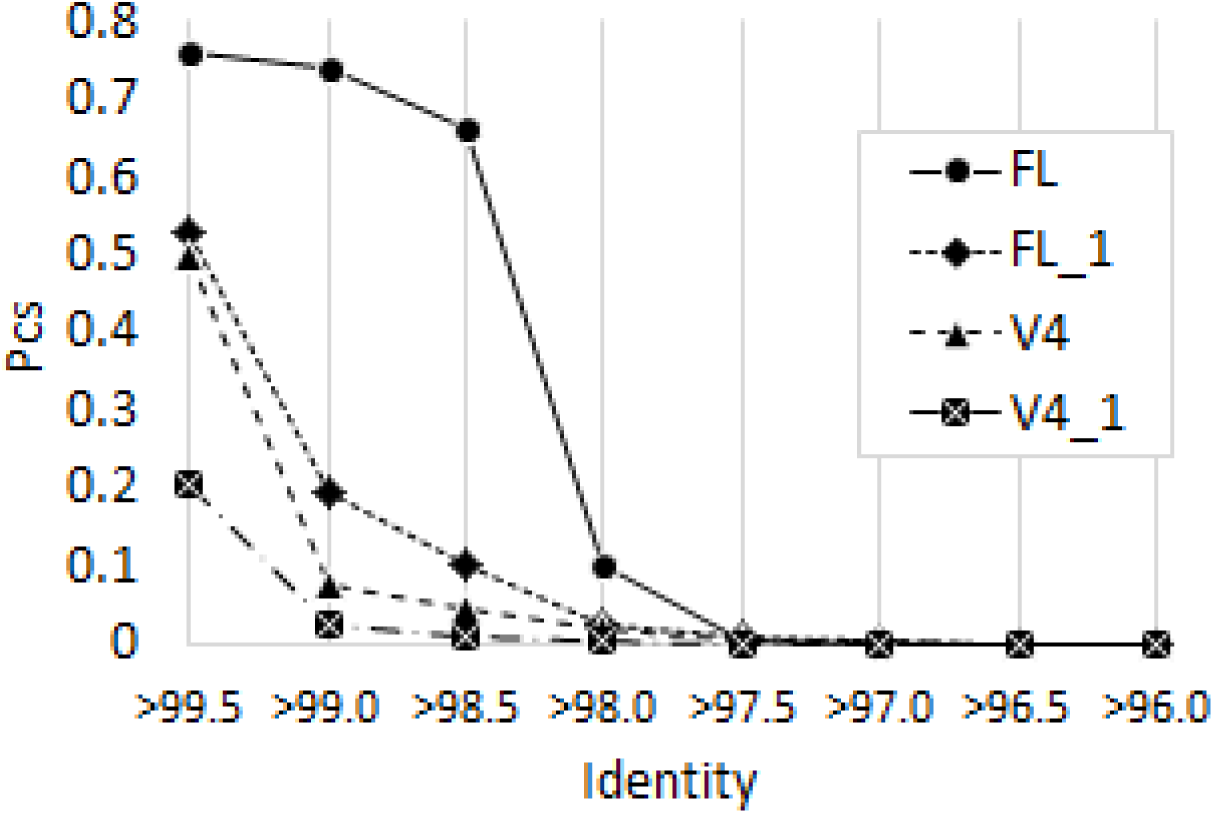
Conspecific probabilities *P*_*sc*_(*d*) for the HiQ databases. FL is HiQFL, FL_1 is HiQFL_1, V4 is HiQV4, V4_1 is HiQV4_1. Identities are binned into intervals of 0.5% so e.g. the *x*-axis label >97% means 97.5%≥*d*>97%.

### Adverse triplets

See Supplementary Files for complete lists of adverse triplets. I found 25 402 triplets in HiQFL_1 with 776/2 512 of species (31%) appearing in at least one triplet. In HiQV4_1, I found 106 576 triplets containing 1 320/2 512 (53%) distinct species, and in soil100 I found 384 triplets containing 25/100 (25%) of the input sequences. This shows that adverse triplets are ubiquitous in the tested datasets and are therefore probably common in practice.

### Mothur alignment errors

A scatterplot of CLUSTALW vs. mothur identities is given in Fig. 2. This shows that mothur systematically underestimates identity of closely-related pairs (>90% identity) compared with CLUSTALW, which constructs alignments by pair-wise dynamic programming (Needleman and Wunsch, 1970). This confirms previous observations (e.g., (Sun *et al*., 2009) that multiple alignments tend to underestimate identity. A manual review revealed that all cases where mothur reported lower identities were due to alignment errors (see Fig. 3 and Supplementary Fig. S1 for an example; all alignments are given in the Supplementary Files). Errors of these types do not occur with pair-wise dynamic programming, which implies that the optimal thresholds reported here may be underestimates for similar clustering methods implemented in other software packages.

**Fig. 2.**
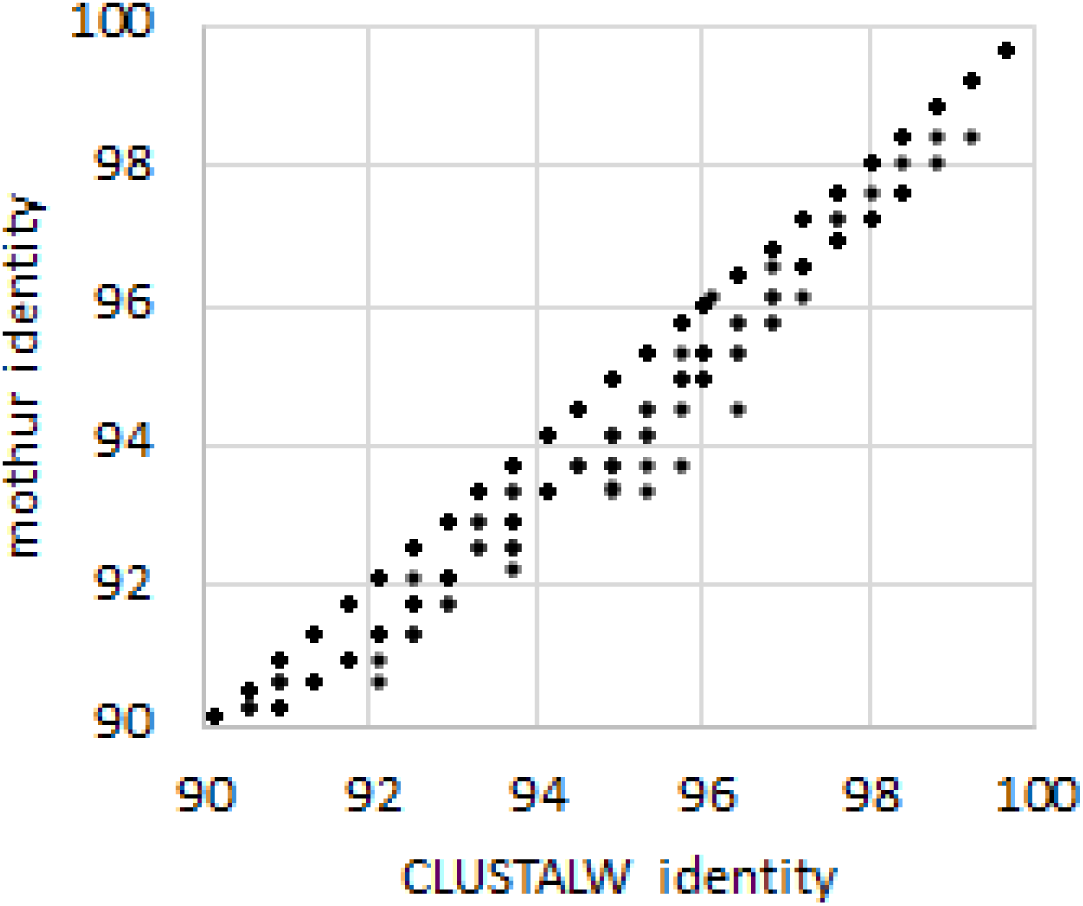
Scatterplot of CLUSTALW vs. identity on the soil100 dataset.

**Fig. 3.**
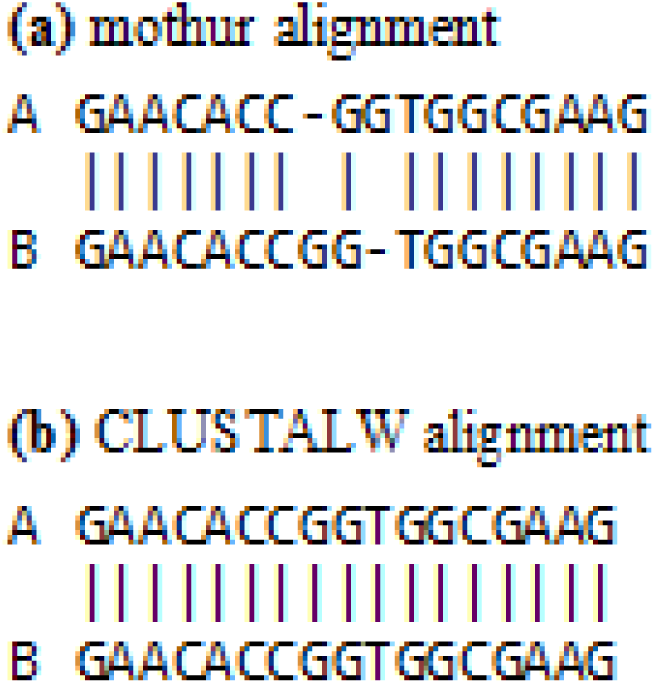
Typical misalignment by mothur. Segment of the alignments by (above) and CLUSTALW (below) for soil.1137 (A) and soil.191 (B); sequences are given in the supplementary data. See Supplementary Fig. S1 for complete alignments. Misalignments of this type do not occur with pair-wise dynamic programming.

## Discussion

### Comments on the MCC _SW_ metric

Recent papers (Schloss and Westcott, 2011; Westcott and Schloss, 2015, 2017) have proposed a variant of Matthews’ Correlation Coefficient (here called *MCC*_SW_) as a definitive accuracy metric for OTUs containing noisy or error-free sequences. Typically, the accuracy of a clustering algorithm is assessed for individual objects (here, sequences) by comparison to categories which have been independently determined (e.g., species). By contrast, *MCC*_SW_ measures accuracy by assuming that true OTUs objectively exist and can be defined indirectly via a binary classification of pairs of sequences from the sequences alone, without considering their biological origin. The standard of truth is based on pair-wise identity as measured by mothur: if a pair is ≥97%, the sequences are asserted to be in the same true OTU; otherwise they are in different true OTUs. However, OTUs by this definition generally do not exist because of adverse triplets. Also, true positives and true negatives by this standard may in fact be biological errors. For example, consider a pair of reads of a chimeric amplicon formed during PCR. They cannot be assigned to a valid biological OTU if they have <97% identity to their parent sequences, but are a true positive if they have >97% identity to each other. Conversely, a pair of paralogs from a single genome should be assigned to the same biological OTU, but are asserted to be a true negative if they have <97% identity. The use of mothur distances as a standard is invalid if misalignments are common, and regardless biases the metric against methods that use different parameters (e.g., gap penalties) or distance measures. The Supplementary Material gives examples where *MCC*_SW_ is undefined due to division by zero and does not give the highest score to the best clusters. Finally, the results presented here show that threshold of 97% is far from optimal as an approximation to species. *MCC*_SW_ is therefore not viable as a benchmark standard of biological accuracy.

### There is no best algorithm, threshold or metric

In the tests reported here, all algorithms achieved comparably high scores by a given quality metric when thresholds were tuned to the input data and metric, showing that no algorithm is intrinsically superior. All metrics were designed to quantify the correspondence between OTUs and species. However, for a given algorithm and dataset different metrics were maximized at different thresholds and thus by different sets of clusters, showing that a single metric cannot definitively quantify accuracy.

### Optimal thresholds are data-dependent

Different thresholds were obtained were obtained on HiQFL compared to HiQFL_1 and on HiQV4 compared to HiQV4_1. These datasets contain the same species with different abundances, and in general it should be expected that the optimal threshold for a given algorithm and quality metric will depend on the segment (full-length, V4 or some other region), composition and abundance distribution of the data. This can be understood from another perspective by considering that the optimal clustering threshold is determined by the underlying conspecific probability function *P*_cs_. The *P*_cs_ function will generally be different for different datasets, as illustrated by the HiQ databases in Fig. 1. Natural communities generally do not resemble reference databases, and are highly variable in their compositions and abundance distributions. Therefore, optimizing a clustering threshold on a given training set cannot reliably predict that it will have high accuracy on novel data.

### The canonical 97% threshold is too low

All thresholds in Table 1 are higher than 97%, especially on the V4 region where all optimal thresholds were >99% with a median of 100%. On full-length sequences, most optimal thresholds (11/20 on HiQFL and 19/20 on HiQFL_1) were >99%. Thus, while keeping the above caveats in mind, it is clear that if the goal of OTUs is to approximate species, then the canonical 97% threshold is far from optimal for all clustering algorithms and should be increased to at least 99%.

### Intra-species variation

There can be “enormous” strain-to-strain variation in gene content within a species (Doolittle and Papke, 2006), causing substantial differences in phenotype. For example, some strains may be pathogenic while others are symbiotic (Ochman, 2001). As a result, OTUs that accurately approximate species will tend to lump distinct phenotypes into a single cluster, and it could therefore be more biologically informative to construct OTUs approximating strains rather than species, raising the question of whether this is achievable in practice. Resolving strains would require a higher identity threshold than species. With the V4 region, optimal thresholds for species are at or very close to 100%, showing that higher resolution is probably not possible in general, though some strains might be resolved for some species. Full-length sequences might enable better strain resolution, as might segments of intermediate length containing two or more hypervariable regions. However, definition and assessment of strain-based OTUs raises new difficulties compared to species. For example, some strains have very similar phenotypes which could reasonably be assigned to the same OTU, while others are substantially different and would preferably be assigned to different OTUs, raising the question of whether such distinctions could be satisfactorily quantified and annotated for parameter training and benchmark testing. Also, while complete genome assemblies are available for multiple strains of many species, yielding a robust set of examples for determining typical levels of intra-species sequence variation, type strains are usually genetically identical rather than naturally occurring subspecies (Dijkshoorn *et al.*, 2000) and little information is therefore available about subspecies sequence variation *in vivo*.

### Zero-radius OTUs (ZOTUs)

If a 100% identity threshold is used, then each distinct sequence defines a separate OTU. I have previously called this a ZOTU (zero-radius OTU) (Edgar, 2016); it has also been called a Sequence Variant (Callahan *et al.*, 2016). I agree with a recent perspective (Callahan *et al.*, 2017) arguing that “improvements in reusability, reproducibility and comprehensiveness are sufficiently great that [ZOTUs] should replace [97%] OTUs as the standard unit of marker-gene analysis and reporting”. ZOTUs achieve the best possible phenotype resolution at the expense of an increased tendency to split species and strains over multiple OTUs. However, some lumping and/or splitting of strains and species is unavoidable at any threshold. With V4 sequences, the results presented here show that ZOTUs achieve a reasonable balance between lumping and splitting of species while 97% OTUs have a strong tendency to lump species together. ZOTUs have the additional advantage of being directly comparable between datasets without re-clustering (i.e., ZOTUs are *stable* as defined by (Rideout *et al.*, 2014)), providing that the same gene segment is compared. With longer sequences, ZOTUs may cause more splitting than lumping, but this is a relatively benign problem which can be addressed by downstream analysis. For example, alpha diversity could be adjusted according to estimated rates of splitting and lumping. ZOTUs of longer sequences may therefore also be preferred over traditional OTUs for their improved ability to discriminate phenotypes.

